# Fluorescence biosensor for real-time interaction dynamics of host proteins with HIV-1 capsid tubes

**DOI:** 10.1101/619841

**Authors:** Derrick Lau, James Walsh, Wang Peng, Vaibhav Shah, Stuart Turville, David Jacques, Till Böcking

**Affiliations:** EMBL Australia Node in Single Molecule Science and ARC Centre of Excellence in Advanced Molecular Imaging, School of Medical Sciences, UNSW Sydney, NSW 2052, Australia; The Kirby Institute, UNSW Sydney, NSW 2052, Australia

**Keywords:** HIV capsid, TIRF, binding, stoichiometry, kinetics, microfluidics

## Abstract

The human immunodeficiency virus 1 (HIV-1) capsid serves as a binding platform for proteins and small molecules from the host cell that regulate various steps in the virus life cycle. However, there are currently no quantitative methods that use assembled capsid lattices for measuring host-pathogen interaction dynamics. Here we developed a single molecule fluorescence biosensor using self-assembled capsid tubes as biorecognition elements and imaged capsid binders using total internal reflection fluorescence microscopy in a microfluidic setup. The method is highly sensitive in its ability to observe and quantify binding, obtain dissociation constants, extract kinetics with an extended application of using more complex analytes that can accelerate characterisation of novel capsid binders.

## Background

HIV-1 establishes infection by integrating its genome into a host chromosome. In order to access the DNA of the target cell, the viral genome must travel across the cytoplasm and enter the nucleus, while avoiding recognition and degradation by the host.^1^ The vehicle responsible for conveying and protecting the genome on this journey is the HIV-1 capsid, a metastable nanocontainer comprising ∼1500 copies of the capsid protein (CA) that assemble into a lattice of hexamers and pentamers to form a fullerene cone. While the capsid physically excludes host macromolecules from accessing the viral genome, its functions are more nuanced than a simple inert protective shell. In addition to avoiding cell autonomous immune responses, the capsid has roles in cytoplasmic trafficking,^2^ reverse transcription,^3^ nuclear entry, and integration site selection.^4^ It must also release the viral genome prior to integration in an enigmatic process called ‘uncoating’ (see Review^5^).

While the CA lattice persists within the target cell, the capsid represents the dominant host-pathogen interface. In this role, the lattice represents a binding platform for the recruitment of numerous host-derived molecules that the virus hijacks to mediate the above processes. Three binding sites on CA have been characterised largely using structural approaches, and each shows affinity towards at least two host cofactors. These sites are: (1) the CypA-binding loop, comprising residues 85–93 which form a flexible structure on the capsid exterior responsible for recruiting cyclophilin A (CypA)^6, 7^ and the nucleoporin Nup358^8^; (2) the FG-binding site, a pocket on the N-terminal domain of CA that binds to phenylalanine-glycine motifs of cleavage and polyadenylation specificity factor subunit 6 (CPSF6)^9, 10^ and a second nucleoporin, Nup153^10^; and (3) the central arginine pore formed by a cluster of five or six symmetry-related copies of Arg18 that recruit deoxyribonucleotides (dNTPs) and inositol hexakisphosphate (IP6).^11, 12^ The CypA-binding loop and FG-binding site are both believed to play roles in nuclear import and evasion of immune recognition. The central arginine pore, on the other hand, is believed to play roles in selective import of dNTPs for reverse transcription and mediating capsid stability through recruitment of polyanions such as IP6. Importantly, the FG-binding site and the arginine pore are only formed in the assembled CA lattice as they contain structural elements that are contributed by multiple adjacent CA monomers. In order to study capsid-binding compounds, it is therefore necessary to use a system that recapitulates the assembled CA lattice.

Current capsid binding assays are usually based on pull-down experiments using capsid as bait from either isolated cores,^13-15^ *in vitro* assembled CA-NC tubes,^16, 17^ or disulfide cross-linked CA A14C/E45C tubes.^18^ These methods are not quantitative and suffer from false positives (due to non-specific interactions) and false negatives (due to weak or transient interactions). Quantitative methods such as isothermal titration calorimetry and surface plasmon resonance are possible using *in vitro* assembled soluble CA A14C/E45C/W184A/M185A hexamers,^19, 20^ but these methods are sample-intense and not all CA interfaces (that would otherwise be present in fully assembled CA lattice for cofactor binding) are represented in the soluble hexamer.

Here we propose to overcome the limitations of existing methods with the design of an interface for capsid binders that has the sensitivity to visualise and quantify binding events at the single molecule level using total internal reflection fluorescence microscopy (TIRFM). The biorecognition element consists of cross-linked and fluorescent CA tubes that are grown on the sensor surface inside a microfluidic device. The biosensor allows accurate measurements of interaction kinetics, affinity and stoichiometry and is compatible with both purified recombinant CA-binders and complex samples such as cell-free expression lysates enabling characterisation of binders that might otherwise be difficult to isolate.

## Materials and methods

### Production and labelling of recombinant proteins

HIV-1 CA (wild type, CA A14C/E45C and CA K158C) was expressed in bacteria and purified as previously described.^21^ CA K158C was labelled as described.^22^ CypA was expressed in bacteria, purified and labelled as described.^22^

### Fluorescence labelling of CPSF6_313-327_

A synthetic peptide from CPSF6 comprising residues 313-327 with a C-terminal cysteine (PVLFPGQPFGQPPLGC, Genscript) was dissolved (1.85 mM) in 50 mM Tris buffer (pH 8) and reacted with Alexa Fluor 488-C5-maleimide (1.85 mM) for 10 min. Thin-layer chromatography (silica gel, methanol:water, 1:1, v/v) showed near quantitative incorporation of the fluorescent dye. CPSF6_313-327_-AF488 was flash frozen and stored at −40 °C. The peptide was diluted to 50 μM with buffer (50 mM Tris, pH 8, 1 mM DTT) prior to use.

### Negative staining electron microscopy

CA assembly mixture (1 M NaCl, overnight, 4°C) was applied to a 200 mesh Cu grid coated with carbon and formvar (Ted Pella) and wicked dry. The sample was stained with a drop of uranyl acetate (2% w/v) and wicked dry immediately. This staining process was repeated three times and the grid was air dried. Micrographs were collected using a FEI Tecnai G2 20 or JEOL1400 transmission electron microscope. Tubes diameters were measured using ImageJ.

### Fabrication of microfluidic devices

An aluminium lithography mould was used to cast poly-dimethylsiloxane microfluidic devices with five parallel flow channels (1000×800×60 μm, L×W×H). The PDMS devices were treated with an air plasma (3 min) and annealed to clean glass coverslips.

### Sensor surface modification and *in situ* assembly of CA tubes

Flow channels were incubated with a solution of PLL(20)-g[3.5]-PEG(2)/PEG(3.4)-biotin(20%) (1 mg/mL, SuSos AG) for 30 min. The channels were washed with water and incubated with a streptavidin (0.2 mg/mL) in blocking buffer (20 mM Tris pH 7.5, 0.4 mM EDTA, 50 mM NaCl, 0.006% NaN_3_, 0.025% Tween 20, 0.2 mg/mL BSA) for at least 20 min. Tubing was inserted into the channel ports and the outlet tubing was connected to a syringe pump operated in withdraw mode at a flow rate of 30 μL/min. Channels were rinsed with 50 μL wash buffer (50 mM Tris, pH 8, 150 mM NaCl), filled with a solution (20 μL, 1:3000 dilution) of biotinylated α-mouse antibody (Jackson ImmunoResearch, 115-066-071) and α-CA antibody (Advanced Biotechnologies, 13-102-100) and again rinsed with wash buffer (50 μL). To grow CA tubes on the sensor surface, a buffer solution (50 mM Tris, pH 8.0, 2 mM DTT, 0.02% NaN_3_) containing CA A14C/E45C (76 µM) and labelled CA K158C (AF488 or AF647, 4 µM) was mixed with high salt buffer (50 mM Tris, pH 8.0, 5 M NaCl) to give a final NaCl concentration of 500 mM, immediately injected into the flow cell and incubated for 30 min. CypA (10 µM) was added to the assembly mixture to reduce tube bundling. The flow channel was then rinsed with wash buffer (50 μL). Before injection of analyte, the flow channel was rinsed with 100 µL of imaging buffer (50 mM Tris, pH 8, 150 mM NaCl, 2 mM Trolox, 2.5 mM protocatechuic acid, m 0.25 U/mL protocatechuate-3,4-dioxygenase).

### Total internal reflection fluorescence (TIRF) microscopy and image analysis

TIRF images were acquired (typical exposure time 10 ms) using an inverted microscope equipped with a Nikon 100× CFI Apochromat TIRF oil immersion objective (1.49 NA), a NicoLase laser system^23^ and two Andor iXon 888 EMCCD cameras. Images were analysed using home-written image analysis software for detection of tubes and extraction of binding/dissociation traces (https://github.com/lilbutsa/JIM-Immobilized-Microscopy-Suite).

### Equilibrium binding analysis to determine K_D_ and stoichiometry of CypA binding to CA tubes

CypA solutions (1–40 μM) were prepared with CypA-AF647 (1 μM) and unlabelled CypA (added to make up the desired final concentration) in imaging buffer. The CypA solution (30 µL) was drawn into the channel (100 µL/min) and incubated for 1 min. TIRF images were recorded in several fields of view (FOV) in both channels using sequential excitation to measure the signals associated with CA tubes (491 nm) and co-localised CypA (639 nm). Subsequently the CypA solution was washed out with imaging buffer (100 μL). Images were analysed to determine the CypA:CA molar ratio reached at equilibrium for each CypA concentration. Binding curves were obtained by plotting the best fit molar binding ratio as a function of CypA concentration. The following equation was used for model fitting to obtain estimates for the dissociation constant and the maximum binding ratio: R(eq) = [CypA] × R(max) / ([CypA] + K_D_), where R(eq) is the CypA:CA ratio at equilibrium for a given CypA concentration, [CypA] is the concentration of CypA, R(max) is the CypA:CA molar ratio at saturation and K_D_ is the dissociation constant.

### Kinetics analysis of CPSF6_313-327_ binding to CA tubes

Imaging buffer containing the analyte (CPSF6_313-327_-AF488, 250 nM) and a free dye as a marker for solution exchange in the flow channel (Alexa Fluor 647, 200 nM) was injected into the flow channel (100 µL/min) and subsequently washed out with imaging buffer (100 µL/min) while recording the CA K158C-AF647 and CPSF6_313-327_-AF488 channels by time-lapse TIRF imaging with a frame rate 1/3 s^−1^ (150 s for wash-in and 300 s for wash-out).

### Biosensor assay of capsid-binding proteins produced by cell-free expression

The open reading frame for CPSF6 (isoform 2) was cloned into a vector for cell free expression of the protein with His_6_-GFP fused to its N-terminus. *In vitro* expression of proteins was initiated by adding plasmid (60 nM) to *Leishmania* extract^24^ supplemented with RNAse OUT (Invitrogen) and the mixture was incubated in the dark at 28 °C for 2.5 hours. Protein concentrations were estimated using single molecule spectroscopy with recombinant eGFP as a standard.^25^ The cell-free expression mixture was diluted with wash buffer (supplemented with 2 mM Trolox, 2.5 mM protocatechuic acid and 0.25 U/mL protocatechuate-3,4-dioxygenase) to a final concentration of 370 nM GFP-tagged protein. This solution (40 µL) was drawn into the flow channel (100 µL/min) and subsequently washed out with imaging buffer (500 µL at 100 µL/min) while recording the CA K158C-AF647 channel and the GFP channel by time-lapse TIRF imaging.

## Results and Discussion

### Biosensor design

The design of the total internal reflection fluorescence (TIRF) imaging-based sensor for measuring the interactions between HIV-1 capsid-binding molecules and self-assembled CA lattices in real time is shown in Figure 1. The sensor interface is assembled on the surface of a glass coverslip that forms the bottom of a microfluidic channel device fabricated by soft lithography in PDMS. Cross-linked and fluorescent CA hexamer tubes are then self-assembled *in situ* and function as the biorecognition elements of the sensor. An analyte solution containing a (putative) capsid-binding species labelled with a different fluorescent dye is flowed through the channel while recording dual colour fluorescence movies using a TIRF microscope equipped with CCD cameras, whereby binding (or dissociation) of analytes can be visualised on individual tubes from the appearance (or disappearance) of fluorescence in the analyte channel. We then use single-molecule analysis of the fluorescence movies to extract kinetic parameters of the interaction. In the following, we first present the development of the sensor interface followed by applications of the sensor to measure binding affinity, stoichiometry and kinetics of host proteins known to interact with the HIV-1 capsid to promote infection.

**Figure 1.**
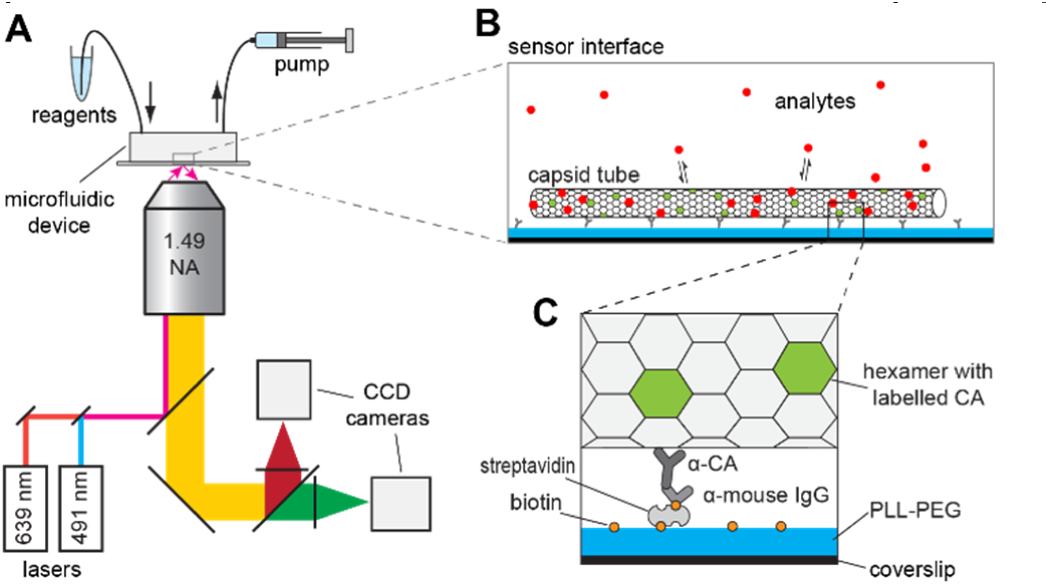
Schematic of the interface design. **A.** The set-up consists of a TIRF microscope for real-time observation of binding events on the sensor surface assembled on a glass coverslip located at the bottom of a PDMS microfluidic device for analyte delivery. **B.** Fluorescent tubes self-assembled from HIV-1 CA serve as the receptor for capsid-binding analytes. Binding and dissociation of analytes is detected by the appearance and disappearance of the analyte signal co-localised with each individual tube. **C.** The sensor surface is modified with an anti-fouling layer and antibodies for immobilisation of fluorescent CA tubes.

### Growth of fluorescent capsid tubes on the sensor surface as biorecognition elements

We used self-assembly of recombinant CA A14C/E45C at high salt concentrations to produce tubular lattices of disulfide cross-linked CA hexamers^26^ as biorecognition elements representing the HIV-1 capsid.^18^ These stabilised tubes were rendered fluorescent by co-assembly with a small fraction of CA K158C reacted with a fluorophore at the engineered cysteine (Supplementary Figure S1 and S2). This co-assembly reaction also yielded long tubes that aggregated into bundles in solution (Figure 2A), a process known to be driven by interactions between hydrophobic residues (primarily located in the CypA loop of CA) on the outer tube surfaces.^27^ Initial attempts to deposit pre-assembled CA tubes onto surfaces failed, presumably because of excessive aggregation into large clusters.

**Figure 2.**
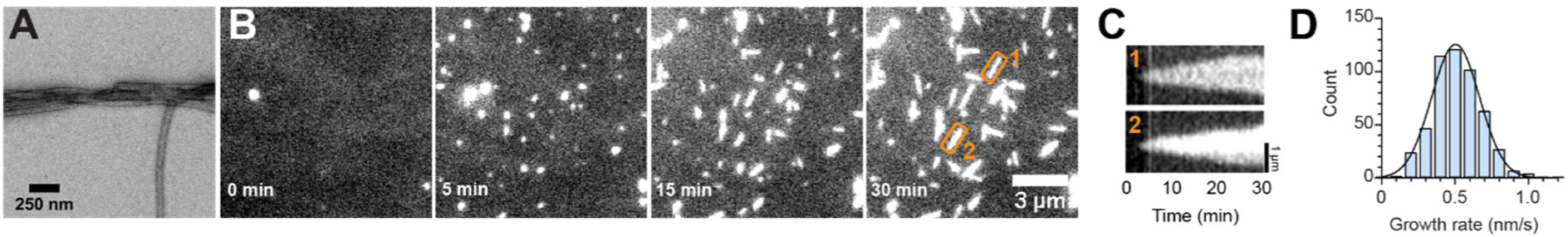
Growth of fluorescent CA tubes on surfaces. **A.** Negative staining EM image of CA tubes assembled from A14C/E45C:K158C-AF568 (72 μM:8 μM). **B.** Time-lapse TIRF images showing the growth of immobilised fluorescent tubes assembled from A14C/E45C:K158C-AF488 (76 μM:4 μM) over 30 min. **C.** Kymographs for two selected tubes highlighted in the last image in panel B. **D.** Histogram of growth rates determined from the slopes of the edges in kymographs (*N* = 515 tubes).

Next, we investigated methods to grow tubes directly on the biosensor surface. The glass surface was first passivated by physisorption of a copolymer of poly(L-lysine) and biotinylated poly(ethylene glycol) (PLL-PEG-biotin) to resist non-specific adsorption of biomolecules from solution (Figure 1C). The PLL-PEG-biotin surface was further modified with streptavidin followed by a biotinylated secondary antibody (anti-mouse IgG F(ab’)_2_ fragment) for capture of a primary mouse antibody directed against the exterior surface of CA.^28^ Injection of an assembly mixture containing CA A14C/E45C and labelled CA K158C into the flow channel followed by incubation (30 min) resulted in the growth of capsid tubes on the modified surface, which appeared as fluorescent lines in the TIRF image (Supplementary Figure S3), which could easily be distinguished from background on the basis of shape and intensity. Fluorescent lines were not observed on surfaces without antibodies (Supplementary Figure S3A) indicating that tube capture was specific. Time-lapse imaging of the assembly reaction on antibody-modified surfaces revealed the tube growth process (Figure 2B). Capsid seeds (presumably small CA assemblies formed in solution) were captured by the antibodies onto the surface where they appeared as bright fluorescent punctae within 5 min of injecting CA assembly solution into the flow channel. Over the next 20 min these seeds grew bidirectionally by addition of CA from solution to the ends of the lattice into ∼ 1 μm long tubes, consistent with previous observations.^28^ The growth kinetics could be determined at the level of single tubes from kymographs generated along the tube axis as shown for two tubes in Figure 2C, whereby the growth rate was given by the slopes of the kymograph edges. Under the conditions used here (500 mM NaCl), capsid tubes ends grew at a rate of 0.52 ± 0.17 nm/s in each direction (Figure 2D).

We observed that the fluorescence intensity along the tubes varied considerably between tubes in the field of view (Supplementary Figure S3B). On the basis of the narrow distribution of tube diameters seen in the negative staining EM images (Figure 3), we speculated that this variation resulted from the growth of multiple tubes bundled together or of multiwalled tubes ^28^ rather than from the growth of individual tubes with varying diameters. It has been shown that wild type CA tube bundling can be minimised by addition of CypA, an 18 kDa globular protein that binds reversibly to the outside of CA tubes.^27^ Assembly of tubes in solution from CA A14C/E45C and labelled CA K158C in the presence of substoichiometric amounts of CypA yielded primarily single tubes with a diameter of 66.9 ± 6.2 nm (slightly larger than tubes assembled in the absence of CypA with a diameter of 57.9 ± 5.8 nm) as determined by negative staining electron microscopy (Figure 3A and B). Some of the tubes appeared to have closed ends. Short tubular or conical structures were also observed.

**Figure 3.**
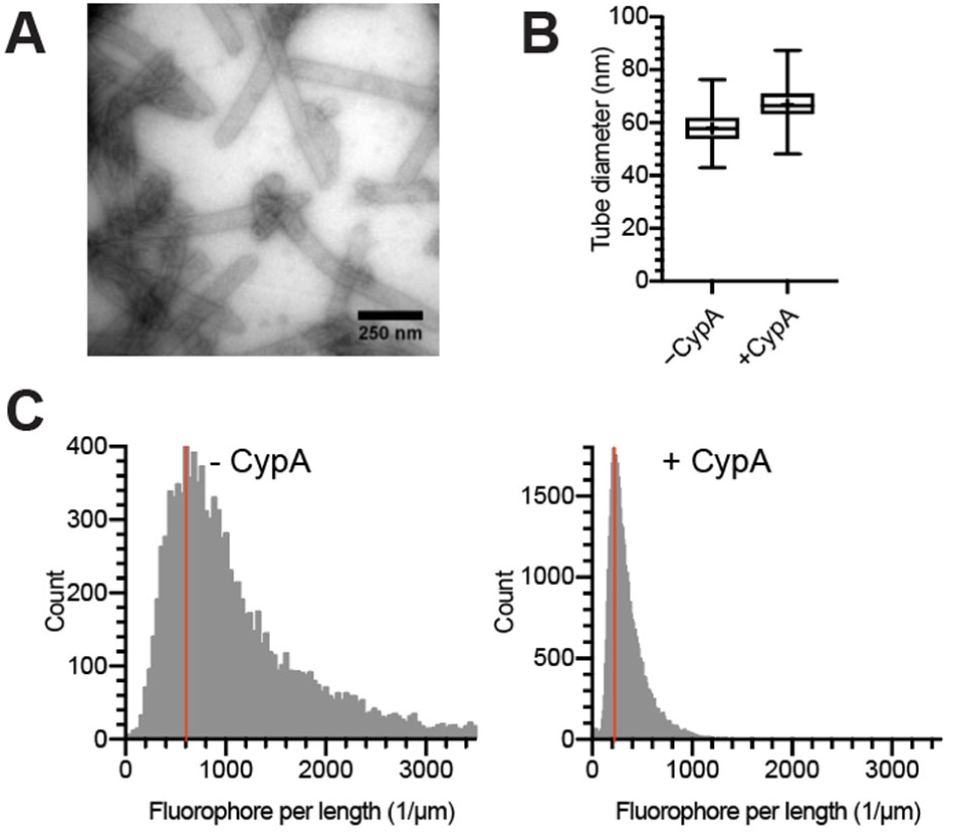
Growth of individual tubes in the presence of CypA. **A.** Negative staining EM image of CA structures assembled from A14C/E45C:K158C-AF488 (76 μM:4 μM) in the presence of CypA (10 μM). **B.** Boxplots of tube diameters measured from EM images from assembly of CA A14C/E45C and labelled K158C in the absence (*N* = 270) and presence (*N* = 235) of CypA. **C.** Histograms of the number of fluorophores per unit length (tubes and bundles) grown on the biosensor surface in the absence (*N* = 10941) and presence of CypA (*N* = 49839). The red line indicates the mean number of fluorophores per unit length.

Next, we tested whether addition of CypA during CA tube growth on surfaces could suppress tube bundling. We grew tubes on surfaces in the absence and presence of CypA and identified fluorescent lines in the TIRF images using an automated image analysis algorithm (Supplementary Figure S4). Analysis of >10000 line-shaped objects showed a linear relationship between the number of fluorophores versus length for both conditions (Supplementary Figure S5), allowing us to represent the data as distributions of fluorophores per unit length. Assembly of CA A14C/E45C and labelled CA K158C in the presence of CypA yielded line-shaped objects with a narrow distribution centered at 223 fluorophores/µm (Figure 3C, right); we assign this linear density of label incorporation to that of single CA tubes grown on the sensor surface. While the majority of objects in this condition were single tubes, the asymmetry of the distribution with a shoulder at higher linear densities indicated that some tube bundles may still be present. In contrast, the distribution of fluorophores per unit length for elongated objects assembled in the absence of CypA was broad with a peak at 604 fluorophores/µm (Figure 3C, left), indicating that most objects assembled in the absence of CypA represent tube bundles with on average 3 tubes. Taken together our data suggest that CypA present during tube growth effectively suppresses tube bundling, allowing us to prepare surfaces with predominantly single tubes. On the basis of the number of fluorophores per unit length measurement for single tubes and the packing of CA molecules in the hexameric lattice (Supplementary Figure S6), we estimated that approximately 1:50– 1:60 CA molecules in the lattice were labelled with a fluorophore, confirming that labelled CA K158C was incorporated less efficiently into the lattice than unlabelled CA. This ratio provided a calibration factor to determine the stoichiometry of analyte binding to capsid tubes in the biosensing experiments described below. Overall our experiments defined parameters for growing individual fluorescent tubes of uniform intensity on passivated surfaces as biorecognition elements, whereby the ability to measure the growth kinetics could also be exploited to test the effects of HIV-1 capsid assembly inhibitors.^29^

### Biosensor assay to measure CypA equilibrium binding curves

In the first application of our biosensor, we used equilibrium binding measurements to determine the dissociation constant (K_D_) and stoichiometry of the CypA-CA complex on CA tubes. CypA is an abundant host cell protein that binds to the CypA binding loop located on the N-terminal domain of CA (Figure 4D). Recruitment of multiple CypA molecules onto the outside of the HIV-1 capsid after host cell entry promotes HIV-1 infection^30^ but the precise binding mode of CypA to assembled CA lattices remains unclear. A solution containing CypA labelled at Cys51 with Alexa Fluor 647 was injected into the flow channel while imaging the fluorescence signals of the cross-linked CA tubes and CypA by dual colour TIRF microscopy. CypA fluorescence appeared rapidly in the CypA channel at locations corresponding to immobilised tubes and reached a constant intensity upon reaching binding equilibrium (Figure 4A). The CypA solution was then washed out and the cycle was repeated for a range of CypA concentrations (1–40 μM). Using automated image analysis we then extracted the number of CA and CypA molecules associated with each tube from the fluorescence intensities in the respective channels (Figure 4B). At any given concentration, the number of bound CypA molecules increased linearly with the number of CA molecules in a tube, i.e. longer tubes bound proportionally more CypA molecules (see example tubes T1, T2 and T3 highlighted in the merged image of Figure 4A and in the inset of Figure 4B). We then calculated the molar binding ratio at each concentration (see Figure 4B for details) to obtain an equilibrium binding curve (Figure 4C). Curve fitting with an equilibrium binding model yielded a K_D_ of 8.0 µM and a stoichiometry of 0.5 CypA per CA at saturation, i.e. half of the CypA binding loops on the CA lattice can be occupied simultaneously as predicted on the basis of structure and simulations.^27^ The K_D_ determined here for the CypA:CA tube interaction was in the range of values for the complex of CypA with unassembled CA previously determined.^4, 31, 32^ Our observations with CypA as an analyte demonstrate the application of the biosensor for accurate affinity measurements, whereby calibration of the response with the intensity of a single molecule also allowed counting of molecules and thus measurement of molar binding ratios. We have recently demonstrated the utility of this analysis by testing the functionality of a proposed secondary CA binding site on CypA.^22^

**Figure 4.**
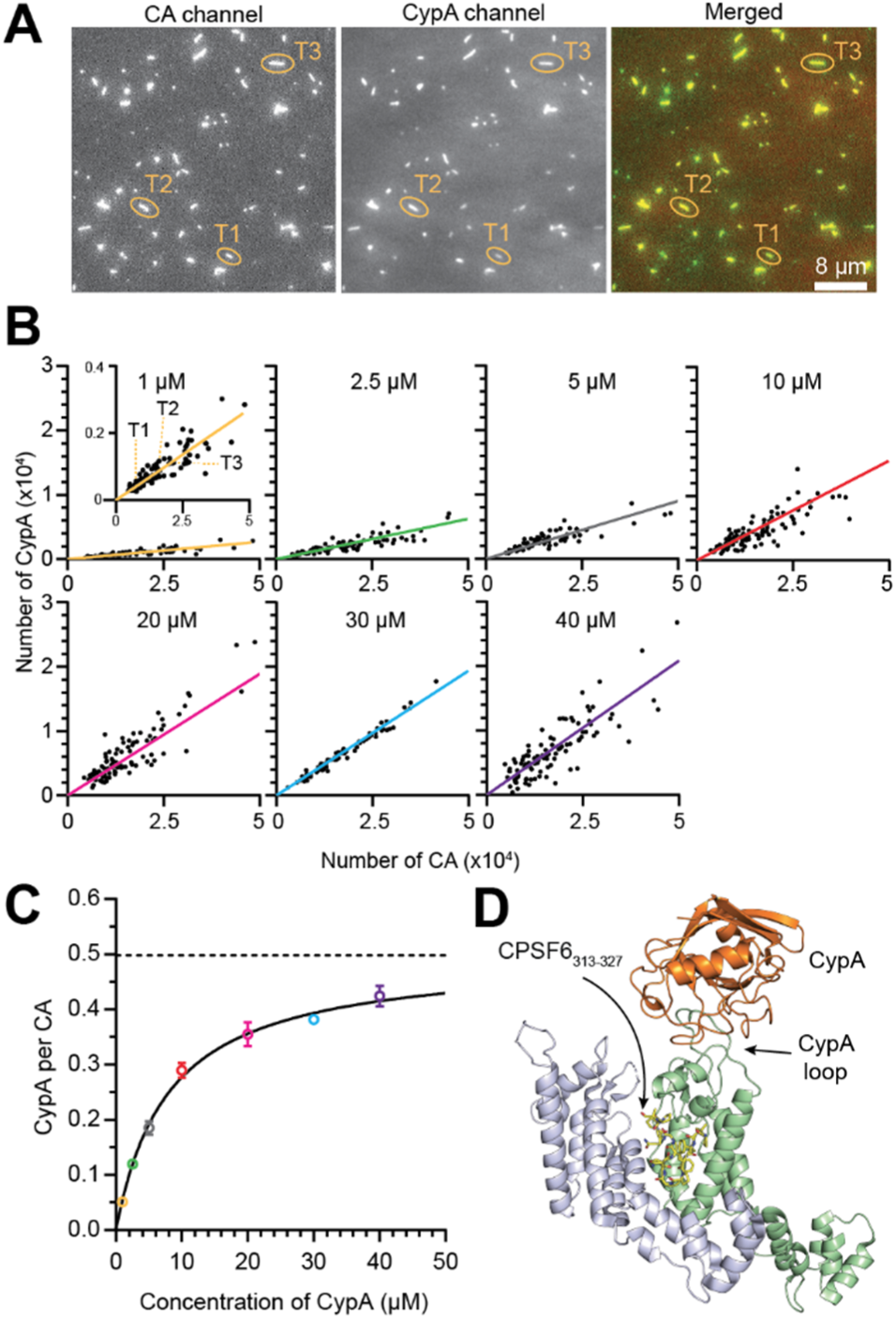
Affinity and stoichiometry of CypA binding to CA tubes. **A.** TIRF images of CA A14C/E45C:K158C-AF488 tubes (left), CypA-AF647 (1 μM) (middle) and overlay of both channels (right). **B.** Scatter plots of the number of CypA molecules versus the number of CA molecules per tube after injection of CypA (1–40 μM); each symbol represents an individual tube. The CypA:CA ratio for each concentration was obtained from the gradient of the line of best fit (coloured line). The inset shows a zoomed in view of the scatter plot recorded at 1 μM CypA-AF647, whereby the data point labelled T1, T2 and T3 correspond to the three tubes highlighted in panel A. **C.** Equilibrium binding curve obtained by plotting the CypA:CA ratio as a function of CypA concentration. Error bars represent mean ± standard deviation of at least three fields of view on the sensor surface. The fit of an equilibrium binding model (black line) estimated a K_D_ of 8.0 μM and CypA:CA molar binding ratio at saturation of 0.5. **D.** Structural model showing two neighboring CA molecules (blue and green) of a hexamer in complex with CypA (orange) and CPSF6_313-327_ peptide (yellow stick model). Generated from PDB 1AK4,^7^ 4U0A^10^ and 3H47.^33^

### Biosensor assay to measure CPSF6 peptide binding and dissociation kinetics

Next, we performed time-resolved biosensor measurements to determine the kinetics of biomolecular interactions on capsid tubes. As an example analyte, we used an Alexa Fluor 488-labelled peptide derived from CPSF6 (residues 313-327), a host cell protein that promotes nuclear import and integration targeting of the viral DNA via interactions with the CA lattice.^34, 35^ CPSF6_313-327_ binds into a pocket formed between adjacent CA molecules in a hexamer (Figure 4D) with low affinity (K_D_ of 50-100 μM).^34, 35^ The analyte was injected into the flow channel at concentration (250 nM) that was >100-fold below the K_D_, i.e. conditions that should lead to less than 1% occupation of available binding sites. To perform kinetics measurements, we used time-lapse dual-colour TIRF microscopy to monitor the appearance and subsequent disappearance of the CPSF6_313-327_-AF488 signal during analyte injection and subsequent washout in the flow channel (Supplementary Figure S7). Using the JIM Immobilized Microscopy Suite of programs (freely available from a GitHub repository, see Materials and Methods) for automated detection of CA tubes and extraction of background-corrected fluorescence signals, we then extracted CPSF6_313-327_-AF488 binding and dissociation traces for each CA tube in a field of view. Example single-tube traces (Figure 5A) and the heatmap of all traces (Figure 5C) showed that both processes followed single exponential profiles. The molar binding ratio at equilibrium reached a value of 0.0049 ± 0.0001 CPSF6_313-327_ per CA (i.e. 0.5% occupation of binding sites) as expected for the low analyte concentration but signal was readily detectable above noise at levels corresponding to 0.1% occupation of binding sites (Figure 5A and Supplementary Figure S7). As a result, kinetic rate constants for binding (k_obs_ = 4.96×10^−2^ s^−1^ at 250 nM) and dissociation (k_off_ = 3.77×10^−2^ s^−1^) could readily be obtained by curve fitting of individual traces (Figure 5A and B) or median traces (Figure 5C).

**Figure 5.**
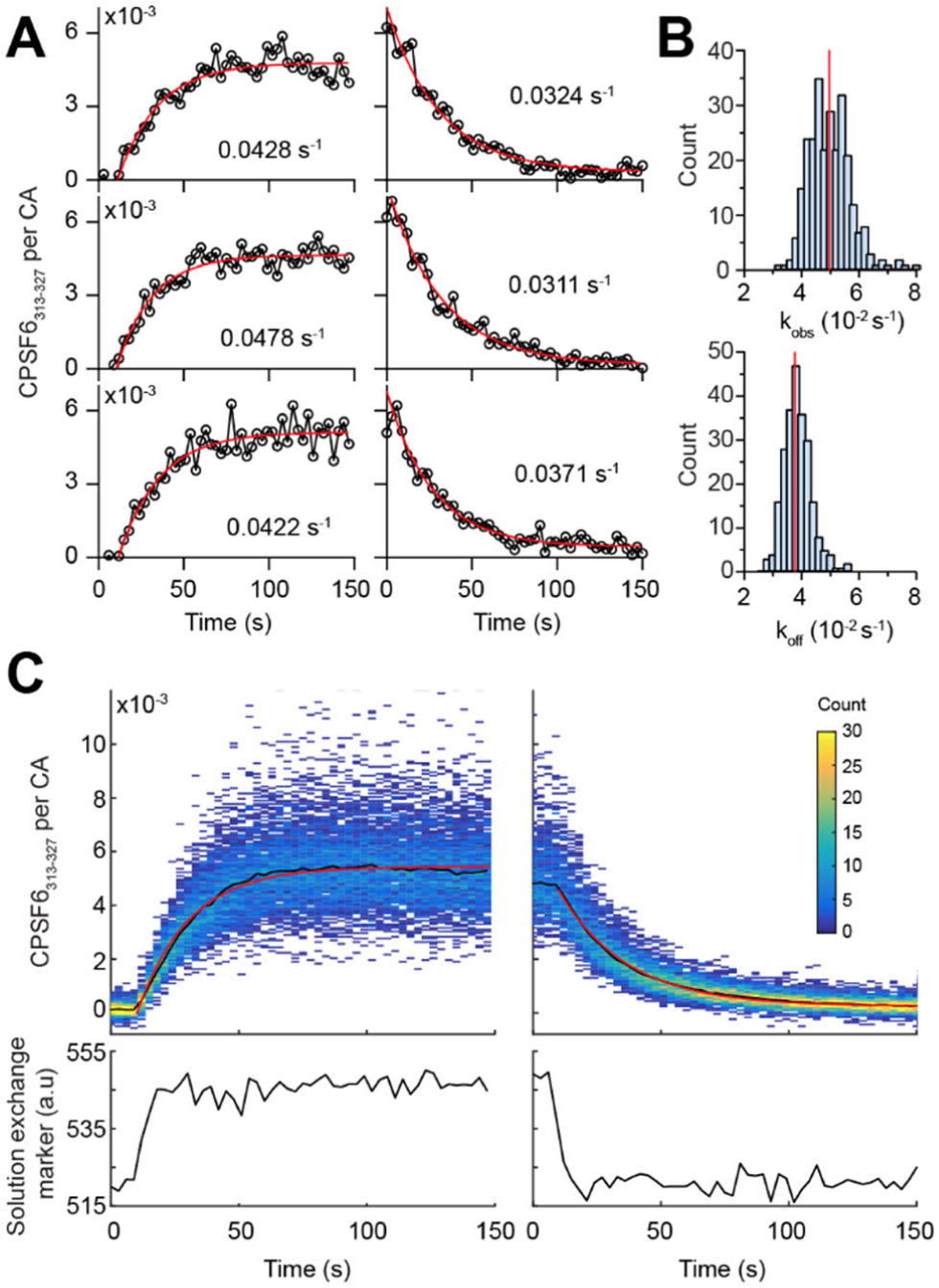
Binding and dissociation kinetics of CPSF6 peptide on CA tubes. **A.** CPSF6_313-327_-AF488 binding (left) and dissociation (right) traces measured on different capsid tubes during CPSF6_313-327_-AF488 injection (250 nM) and wash-out, respectively. The rate constants (*k*_obs_ for binding and *k*_off_ for dissociation traces) were determined from least-squared single exponential fitting (red line) of the fluorescence traces. **B.** Histograms of values obtained for *k*_obs_ (top) from fitting individual binding traces (*N* = 272 tubes) and for *k*_off_ from fitting individual dissociation traces (*N* = 238 tubes). The red lines indicate values from the fitting of median traces in A. **C.** Heatmaps (top) showing the CPSF6_313-327_:CA ratio for all tubes in the field of view during CPSF6_313-327_-AF488 injection (250 nM, left) and washout (right). The median CPSF6_313-327_-AF488 traces (black line) were fitted with exponential functions (red line) to give rate constants of *k*_obs_ = 4.96×10^−2^ s^−1^ and *k*_off_ = 3.77×10^−2^ s^−1^, respectively. The signal of a marker for solution exchange in the flow channel is shown in the plots at the bottom.

### Binding analysis of CPSF6 produced by cell-free protein expression

Finally, we tested the compatibility of the biosensor with GFP-fusion proteins produced by cell-free expression, a rapid method for producing recombinant proteins in microtiter plates for screening protein-protein interactions.^24, 36^ Expression of full length CPSF6 fused at its N-terminus to GFP (GFP-CPSF6) in *Leishmania* extract yielded a mixture of different oligomeric states (Supplementary Figure S8), as previously observed for N-terminally tagged CPSF6_1-358_ expressed in mammalian cells.^37^ In contrast, expression of GFP as a control protein yielded monomers (Supplementary Figure S8). We then injected cell-free expressed proteins without further purification (adjusted to a concentration of 370 nM) into the flow channel of the biosensor to detect the equilibrium binding level of the proteins on CA tubes. As expected, GFP did not bind to sensor interface (Figure 6A and B). GFP-CPSF6 bound strongly to the tubes (Figure 6A) and reached a level of 0.20 ± 0.03 molecules per CA (Figure 6B), which decayed slowly after wash-out (Figure 6C), while the signal of CPSF6_313-327_-AF488 reached a 30-fold lower level and decayed rapidly (Figure 6B and C). These observations are consistent with a strengthening of the CPSF6-capsid interaction through avidity and with oligomerization playing an important role in recognizing viral capsids as suggested before.^37^ Further experiments are needed to identify the physiologically relevant oligomeric species and dissect their effects on the functions of CPSF6 in the viral replication cycle.

**Figure 6.**
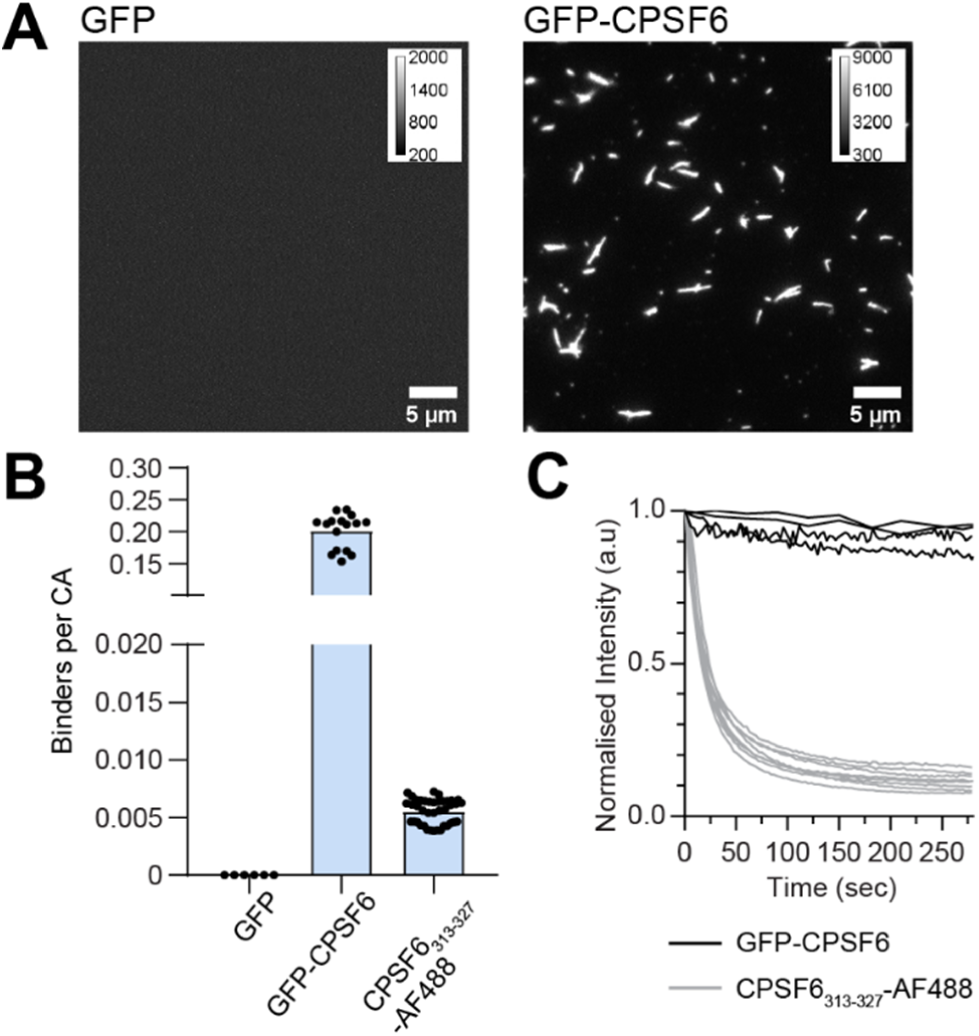
Binding analysis of proteins produced by cell-free expression. **A.** TIRF images of the analyte channel after injection of GFP (left) and GFP-CPSF6 (right). **B.** Bar graph of the molar binding ratio between analyte to CA. Each symbol represents the median value from a field of view taken from at least two different flow channels. **C.** Dissociation traces of GFP-CPSF6 and CPSF6_313-327_-AF488 during wash-out. Each curve represents a separate experiment.

## Conclusions

Our TIRF imaging-based biosensor is ideal for time-resolved and quantitative interaction measurements between capsid-binding molecules and HIV-1 CA tubes to obtain kinetic rate constants, binding affinity and stoichiometry. The key advantage of this system is its stable and reusable interface which allows robust detection of analyte binding at levels corresponding to single molecules per viral capsid (i.e. one analyte per 1500 CA molecules). The appearance of the capsid tubes as lines in the fluorescence image facilitates detection and extraction of traces using automated image analysis. Tubular CA lattices are commonly used in binding studies as surrogates for the capsid^16, 18, 38, 39^ but lack CA pentamers and the variable curvature of the native conical capsid. Combining our biosensor as a tool for routine measurements with more specialised approaches using virus-like particles^32^ should overcome these limitations.

The ability to monitor binding dynamics of different species labelled with distinct fluorophores at the level of single tubes with high sensitivity and temporal resolution opens the door to resolving binding cooperativity and competition, similar to TIRF-based measurements of other linear biopolymers^40-43^. The compatibility of the biosensor with complex samples and low reagent usage could facilitate analysis of proteins in mammalian cell lysates that are otherwise difficult to produce (such as CPSF6^37^ and TRIM5α^39, 44^) and rapid screening of potential new interactors of the HIV-1 capsid produced by cell-free expression. Proof-of-principle measurements demonstrated the utility of the HIV-1 capsid biosensor for testing binding modes of host proteins involved in promoting HIV-1 infection, suggesting a role of oligomerization for CPSF6 binding. The underlying design can conceptually be extended to other viruses^45-48^ for discovery and characterization of proteins, small molecules and drugs that interact with viral capsids to facilitate or inhibit gene delivery and viral replication.

## Supporting information

Supporting information (PDF) includes Supporting Materials and Methods, Supporting Results and Supporting Figures (SDS PAGE analysis of CA K158C fluorescence labelling kinetics and assembly efficiency; Negative staining EM of CA tubes with and without labelled CA K158C; TIRF images of CA tubes grown on surfaces with and without antibody; Output of analysis software developed for automated detection and masking of capsid tubes in fluorescence images; Heatmaps of the number of fluorophores as a function of length for elongated structures grown on the sensor surface in the absence and presence of CypA; Schematic of CA tube geometry; TIRF images of CPSF6_313-327_-AF488 binding and dissociation; Single molecule spectroscopy of proteins produced by cell-free expression using *Leishmania* extract).

## Author contributions

Derrick Lau – investigation (protein production/labelling/assembly, assay development, TIRFM binding assay with CypA and CPSF6), data analysis, writing – original draft; James Walsh – software, data analysis, writing–review and editing; Wang Peng – investigation (TIRFM binding assay with CypA), data analysis, writing–review and editing; Vaibhav Shah – supervision, writing–review and editing; Stuart Turville – supervision; writing–review and editing; David Jacques – supervision, writing – original draft; Till Böcking – conceptualisation, supervision, writing–original draft.

## Acknowledgments

We thank Owen Pornillos (University of Virginia) for supplying expression constructs for CA mutants; Alex Ma (UNSW) for help with molecular biology; Yann Gambin and Emma Sierecki (UNSW) for supplying *Leishmania* extract; Philip Nicovich (current address: Allen Institute for Brain Science, Seattle, USA) for building the TIRF microscope; Andrew Tuckwell for providing eGFP for SMS calibration. EM data was acquired using facilities at the Electron Microscope Unit at UNSW.

---

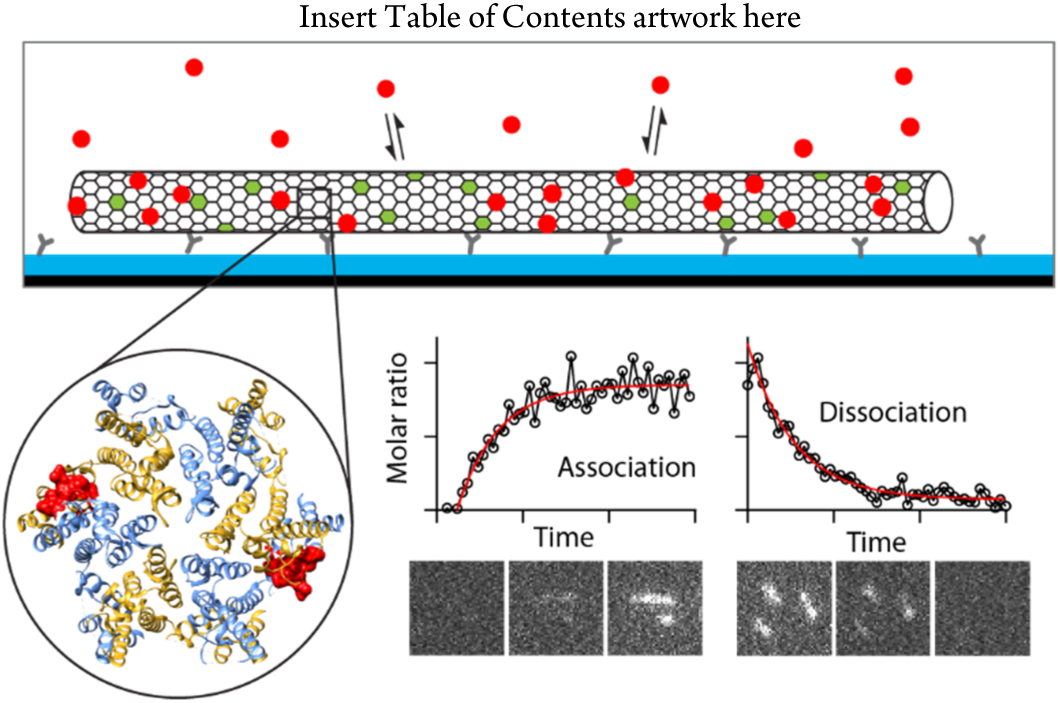

