## Supplement_information for "Fluorescence biosensor for real-time interaction dynamics of host proteins with HIV-1 capsid tubes"

### Supporting Materials and Methods

**Purification of recombinant CA.** Constructs for expression of untagged wild type CA and CA A14C/E45C in a pET11a vector were a gift from Owen Pornillos. The construct for expression of CA K158C was generated from the plasmid containing wild type CA using site directed mutagenesis. CA was expressed in *E. coli* BL21 Rosetta (DE3, pLysS RARE) grown in Luria-Bertani medium supplemented with ampicillin (100 µg/mL) and chloramphenicol (34 µg/mL) at 37 °C with shaking. Protein expression was induced by adding IPTG to a final concentration of 0.4 mM when the optical density at 600 nm reached 0.5 and allowed to proceed at 20 °C for 16 hours with shaking. Cells were harvested by centrifugation, resuspended in lysis buffer (50 mM Tris, pH 8, 2 mM DTT, 0.02% $NaN_3$, 1 mg/mL lysozyme, Complete protease inhibitor) and lysed by sonication. For purification of CA A14C/E45C, the lysis buffer contained 200 mM of 2-mercaptoethanol to prevent disulfide crosslinking. For purification, the cell lysate was clarified by centrifugation (43150 g, 1 hour, 4°C). CA assembly was induced by adding solid NaCl (2.5 M) and the solution was incubated for 15 min on ice and then pelleted (3900 g, 45 min, 4°C). The supernatant was removed and the pellet was redissolved in buffer A (50 mM Tris, pH 8, 2 mM DTT, 0.02% $NaN_3$) to induce disassembly of CA. CA assembly was repeated by adding NaCl (2.5 M), the assembled protein was pelleted and redissolved in buffer A as above. The solution was then filtered (0.22 µm) and CA was further purified by subtractive anion exchange chromatography using a HiTrap Q HP column (GE Healthcare, 17115301). The column was washed with 5 column volumes (CV) of buffer A, followed by 5 CV of buffer B (buffer A containing 1 M NaCl) and then equilibrated with 5 CV of buffer A at 5 mL/min before injecting the sample at 1 mL/min. CA was collected in the flowthrough and fractions containing CA were identified using reducing SDS-PAGE. Protein concentrations were determined using the Bradford assay (Thermo Fisher Scientific, 23236) with BSA as a standard. CA protein solutions in buffer A were concentrated to a concentration of ≥ 150 µM (3.8 mg/mL) using centrifugal ultrafiltration devices (Amicon Ultra-15, 10 MWCO, Merck, UFC901008), snap frozen in liquid nitrogen and stored at -80 °C.

**Fluorescence labelling CA K158C.** CA K158C tubes were assembled (4 °C, 15 min), pelleted and washed three times with labelling buffer (50 mM, pH 8, 0.1 mM TCEP, 0.02% $NaN_3$, 2.5 M NaCl) to remove DTT as described above. A maleimide derivative of a fluorescent dye (Alexa Fluor 488-C5-maleimide [A10254], Alexa Fluor 568-C5-maleimide [A20341] or Alexa Fluor 647-C2-maleimide [A20347]; Thermo Fisher Scientific) was added to assembled tubes at 2-fold molar excess and reacted for 2 min at room temperature before addition of 2-mercaptoethanol (25 mM) to quench unreacted dye. The labelled tubes were collected by centrifugation (18000 g, 10 min, 4°C) and resuspended in labelling buffer to remove excess dye. This washing process was repeated three times. Then the tubes were pelleted and resuspended in buffer A without NaCl (50 mM Tris, pH 8, 0.1 mM TCEP, 0.02% $NaN_3$) and incubated overnight at 4 °C to induce tube disassembly. Finally the solution was centrifuged to remove aggregates and the supernatant containing labelled CA K158C was recovered. The degree of labelling determined using UV-visible spectroscopy was typically ≥85%. Proteins were snap frozen in liquid nitrogen and stored at -40 °C.

**Fluorescence labelling of recombinant CypA.** Purification of untagged recombinant CypA was carried out as previously described.[1] Recombinant CypA was quantitatively labelled using a 2-fold molar excess of Alexa Fluor 647-C2-maleimide in PBS (pH 7.4) containing 0.1 mM TCEP at room temperature for 10 min and the reaction was quenched by adding DTT. Excess dye was removed using Zeba desalting columns (Thermo Fisher Scientific, 89882) equilibrated with CypA storage buffer (50 mM Tris, pH 7.9, 1 mM DTT, 20% glycerol) and Alexa Fluor 647 labelled CypA (CypA-AF647) was flash frozen in liquid nitrogen for storage at -40 °C.

**Assembly of CA tubes in solution.** CA stock solution in buffer A were brought to the desired concentration with Tris buffer (50 mM, pH 8) and assembly was achieved by addition of 1 M NaCl followed by overnight incubation at 4 °C. The assembly mixtures contained the following CA proteins (final concentrations): (1) wild type tubes (80 µM WT); (2) labelled tubes (72 µM WT + 8 µM K158C-AF488); (3) cross-linked tubes (80 µM A14C/E45C); (4) cross-linked and fluorescent tubes (72 µM A14C/E45C + 8 µM K158C-AF488 [or AF568]); (5) cross-linked and fluorescent tubes assembled in the presence of CypA (76 µM A14C/E45C + 4 µM K158C-AF488 + 10 µM CypA).

**Biochemical characterization of CA tube assembly efficiency.** Assembly efficiency was determined by collecting tubes by centrifugation (18000 g, 10 min, 4 °C) and analysing supernatant and pellet fractions by reducing SDS-PAGE with fluorescence detection and Coomassie staining followed by densitometry (ImageJ).

**TIRF microscopy image analysis.** The fluorescence intensity of single fluorophores was determined from the quantal bleaching step in intensity traces of labelled proteins adsorbed sparsely to the coverslip surface and imaged continuously. The number of labelled molecules (labelled CA K158C or analyte protein) associated with a tube was determined by dividing the total intensity of the tube in the channel of interest by the corresponding single-molecule intensity. The fraction of labelled CA K158C protein incorporated into the lattice of cross-linked tubes was estimated by relating the mean number of fluorescent labels per unit length (Figure 3C) to the total number of CA proteins per unit length (calculated using the tube diameter determined by negative staining EM and geometric considerations of the hexameric lattice, see Supplementary Figure S6). This fraction of label incorporation was then used to calculate the total number of CA proteins in a tube. Finally, the analyte:CA molar ratio was determined by dividing the number of analyte molecules associated with a tube by the total number of CA proteins in the same tube.

**Imaging of tube growth on surfaces.** Tube growth was imaged by time-lapse TIRF microscopy at a frame rate of 1/30 s-1 (5 mW laser power, 10 ms exposure time). Kymographs were generated from time lapse movies using the MultipleKymograph plugin in ImageJ. Each tube typically grew bidirectionally resulting in a triangle shaped kymograph where the apex is the seed. The growth rates were estimated by determining the slopes of the sides of the triangle.

**Supporting Results**

**Site-specific labelling of CA and self-assembly of fluorescent CA tubes.** To obtain fluorescent CA for assembly of capsid tubes on the sensor surface, we produced recombinant CA with the lysine at position 158 mutated to cysteine for labelling with a dye using maleimide click chemistry (Supplementary Figure S1). The engineered cysteine is exposed on a loop located on the inside of the assembled capsid where it is away from interfaces involved in lattice contacts or analyte binding. We expected the native cysteines (C198 and C218) to be poorly accessible for modification in the context of the assembled CA lattice. A labelling time course of assembled CA showed that a maleimide derivative of Alexa Fluor 488 (AF488) reacted rapidly with CA K158C but not wild type CA, while additional slow incorporation of label was observed in both cases (Supplementary Figure 1B and C). These observations suggest that the engineered cysteine residue was highly reactive while non-specific labelling at other residues was slow.

Next, we tested whether CA K158C-AF488 could co-assemble with wild type CA to form larger structures. Self-assembly of CA was induced by high salt and assembled particles were collected by centrifugation. Analysis of supernatant and pellet fractions by SDS-PAGE with Coomassie staining (Supplementary Figure S1D) followed by densitometry (Supplementary Figure S1E) showed that the fraction of pelletable CA in a mixture containing wild type CA and CA K158C-AF488 at a molar ratio of 9:1 (24.4 ± 9.8%, mean ± standard deviation) was similar to wild type CA (31.6 ± 8.9%), confirming that CA assembly was efficient in the presence of the labelled species. Fluorescence was present in the pellet (Figure 2D) but the fraction of CA K158C-AF488 in the pellet was only about half (13.9 ± 8.3%) compared to that of wild type CA. We conclude that overall CA assembly is efficient in the co-assembly mixture but the labelled species is incorporated with about half the yield of wild type CA.

Wild type CA self-assembles into long tubes at high salt but mutations in CA can give rise to other architectures such as cones or spheres.[2] Thus, we used negative staining electron microscopy to investigate the effect of labelled CA K158C on the structure of the assembled lattices. Co-assembly of wild type CA and CA K158C-AF488 at a molar ratio of 9:1 yielded long tubes with a diameter of 45 ± 8.6 nm, indistinguishable from wild type CA tubes with a diameter of 46 ± 6.7 nm (Supplementary Figure S2). Finally, we investigated the ultrastructure of cross-linked CA tubes assembled in the absence and presence of labelled CA. Wild type CA tubes are only stable at high salt, conditions which are not ideal for biosensing. To enable biosensor operation at low salt, we decided to use CA A14C/E45C tubes that are stabilized by disulfide cross-links formed between adjacent CA in each hexamer of the lattice.[2] The diameter of CA A14C/E45C tubes was essentially the same when assembled in the absence (63 ± 6.7 nm, mean ± standard deviation) or presence (58 ± 5.8 nm) of labelled CA K158C (Supplementary Figure S2), but both conditions yielded significantly wider tubes in comparison to wild type CA tubes (see above). Taken together our data show that CA K158C site-specifically labelled at the engineered cysteine is readily incorporated into CA tubes without major effects on tube morphology.

**Supporting Figures**

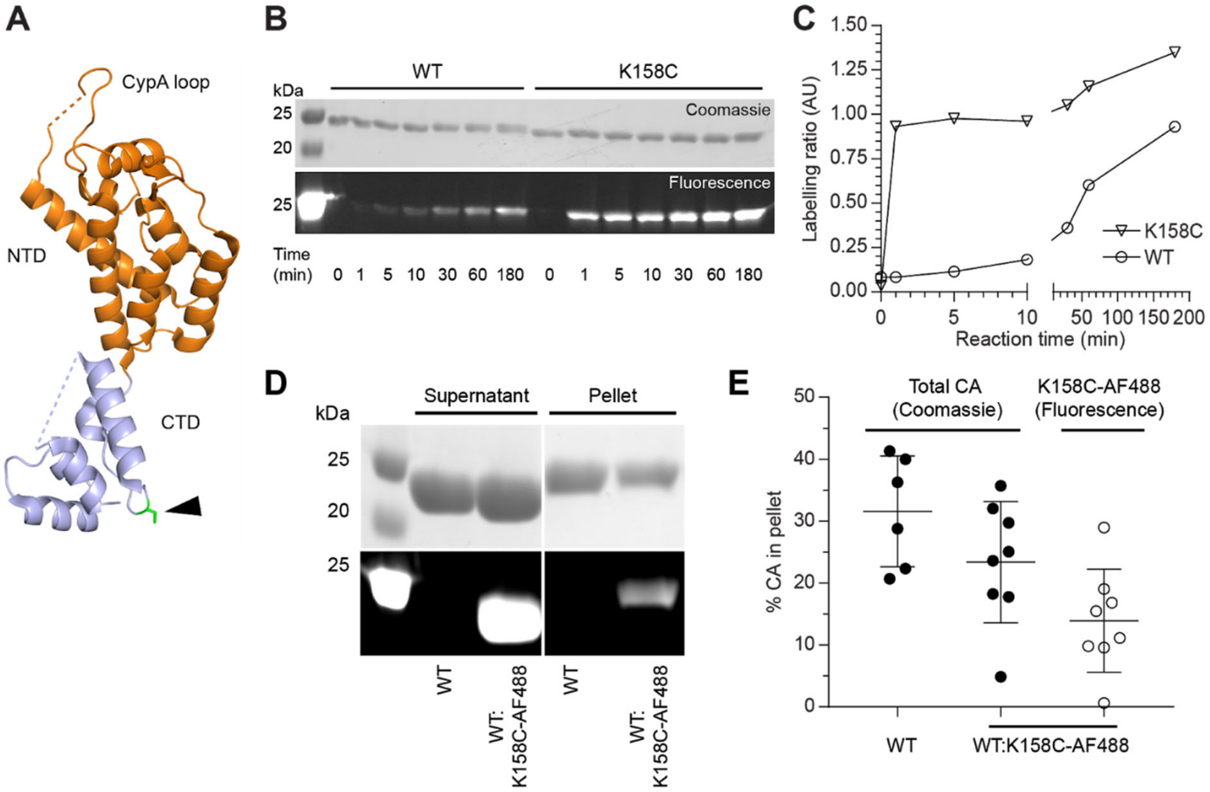


**Supplementary Figure S1. Fluorescence labelling kinetics and assembly efficiency of CA K158C. A.** Ribbon diagram of CA showing the cyclophilin A binding loop (CypA loop), N-terminal domain (orange, NTD) and C-terminal domain (cream blue, CTD). The engineered cysteine residue was modified at position 158 (green/black arrow) from the CA crystal structure (PDB 5HGN).[3] **B/C.** Labelling kinetics of CA (WT or K158C) with AF488-maleimide. CA was assembled into tubes and reacted with AF488-maleimide. The labelling reaction was quenched by addition of 2-mercaptoethanol at the indicated time points and label incorporation was measured by SDS-PAGE with Coomassie staining and fluorescence detection (B). The labelling ratio as a function of time was determined from the band intensities (C). **D/E.** Efficiency of CA WT assembly and of co-assembly of WT:K158C-AF488 (9:1, mol/mol). CA assembly reactions were centrifuged and supernatant (unassembled CA) and pellet (assembled CA) fractions were analysed by SDS PAGE analysis with Coomassie staining and fluorescence detection (D). The overall assembly efficiency and the assembly efficiency of CA K158C-AF488 were determined from the Coomassie and fluorescence signals respectively. Each symbol represents independent assembly reaction with error bars representing mean ± standard deviation.

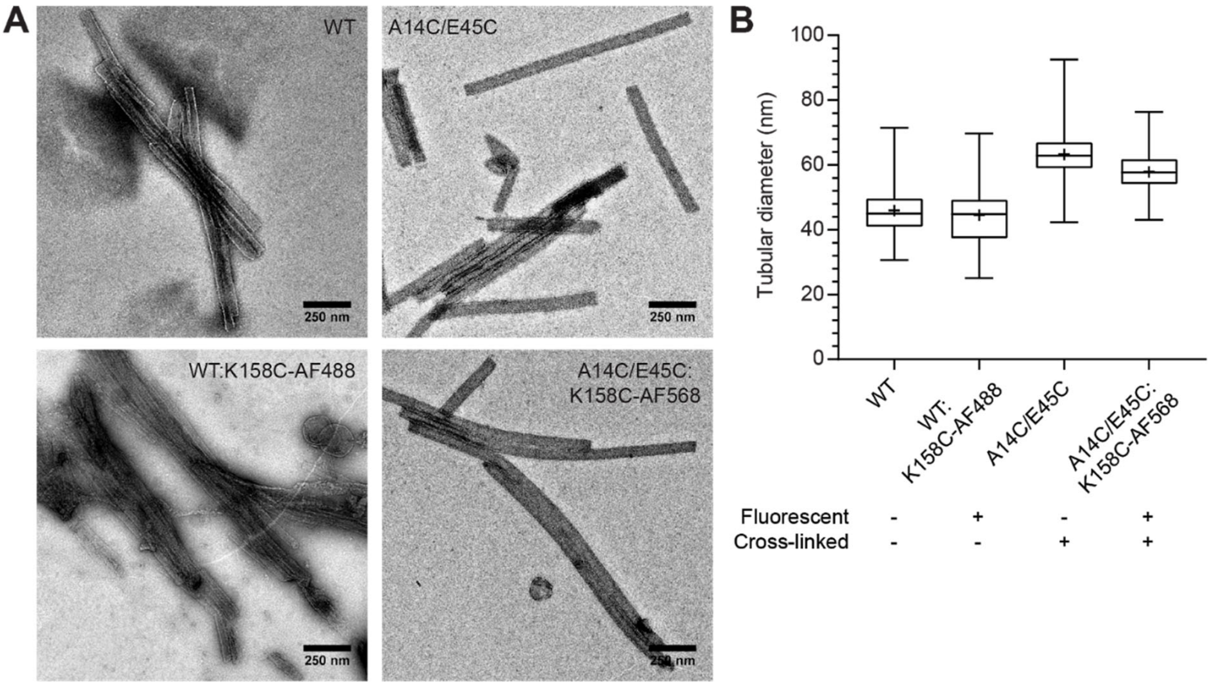


**Supplementary Figure S2. Morphology of CA tubes containing labelled CA K158C. A.** Negative staining EM images of CA tubes self-assembled from reaction mixtures containing WT (control), WT:K158C-AF488 (9:1, mol/mol) for labelling, A14C/E45C for cross-linking and A14C/E45C:K158C-AF568 (9:1, mol/mol) for cross-linking and labelling. **B.** Boxplots of tube diameters measured from the EM images; WT (*N* = 263), WT:K158C-AF488 (*N* = 84), A14C/E45C (*N* = 475), A14C/E45C:K158C-AF568 (*N* = 270).

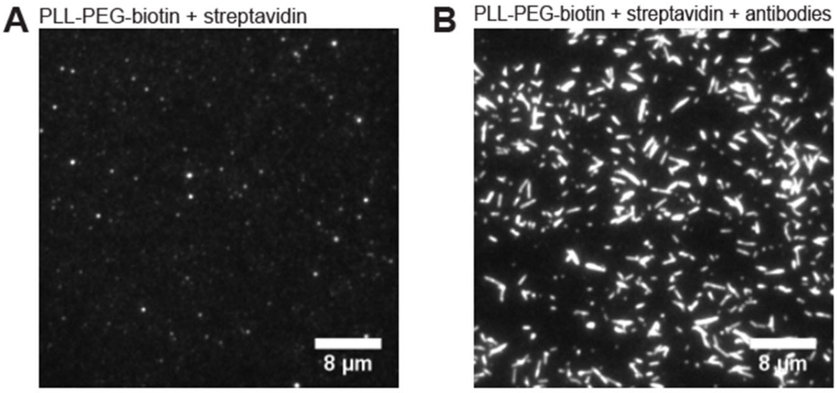


**Supplementary Figure S3. Specific capture of CA for growing tubes on surfaces.** TIRF images of PLL-PEG-modified sensor surfaces without (A) and with (B) antibodies directed against CA recorded 30 min after injection of a CA assembly reaction mixture into the flow channel.

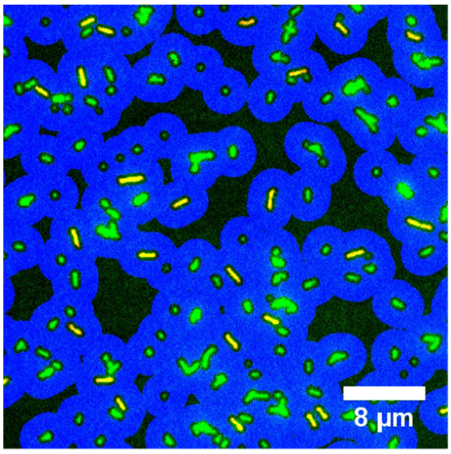


**Supplementary Figure S4. Output of analysis software developed for automated detection and masking of capsid tubes in fluorescence images.** Objects were identified on the basis of intensity and shape as straight lines with uniform intensity corresponding to tubes and tube bundles (highlighted in yellow) or overlapping/irregular/short objects (highlighted in green). The blue masks identify background regions surrounding fluorescent objects and were used for background correction of the fluorescence signal associated with tubes. Data from tubes/tube bundles (yellow objects) were used to create the plots of fluorescence intensity as a function of tube length in Figure 3C.

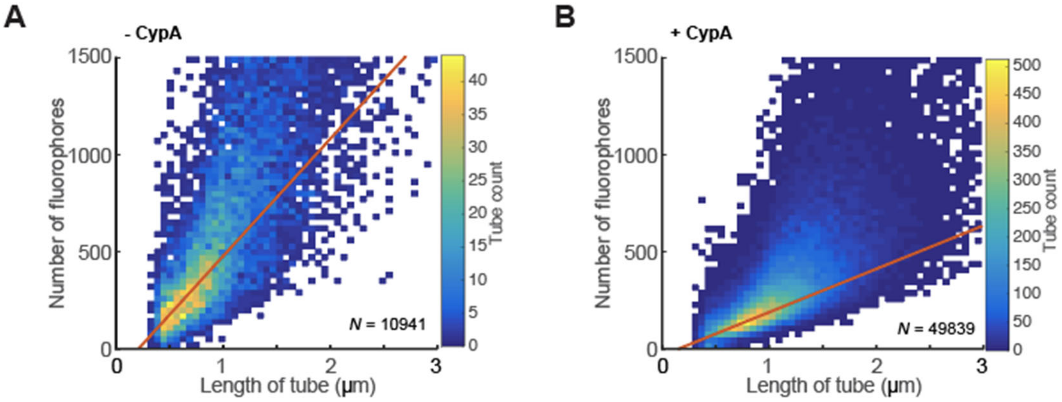


**Supplementary Figure S5. Effect of CypA on bundling of tubes on surfaces.** Heatmaps showing the number of fluorophores as a function of length for elongated structures (individual tubes and tube bundles) grown from A14C/E45C:K158C-AF488 on the sensor surface in the absence (A) ($N$ = 10941 tubes) and presence (B) ($N$ = 49839 tubes) of CypA; combined data from ≥ 3 flow cells. The number of fluorophores for each object was obtained by dividing its total fluorescence intensity by the absolute intensity of a single molecule. The slope of a least-squares fit (red line) gives the following estimates for the number of fluorophores per unit length: assembly without CypA, 604 fluorophores/µm; assembly with CypA, 223 fluorophores/µm.

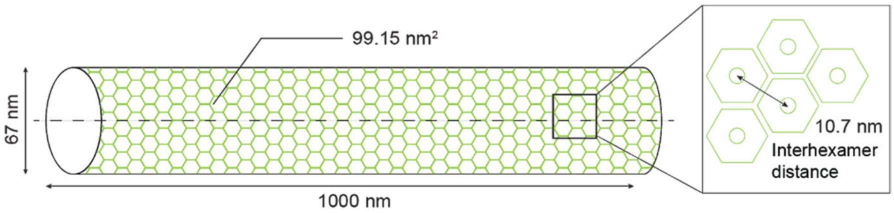


**Supplementary Figure S6. Schematic of CA tube geometry.** Capsid tubes are composed of arrays of CA hexamers with an interhexamer distance of 10.7 nm,[4] corresponding to a surface area of 99.15 nm$^2$ per hexamer. A tube with a diameter of 67 nm contains 12737 CA molecules per µm of length.

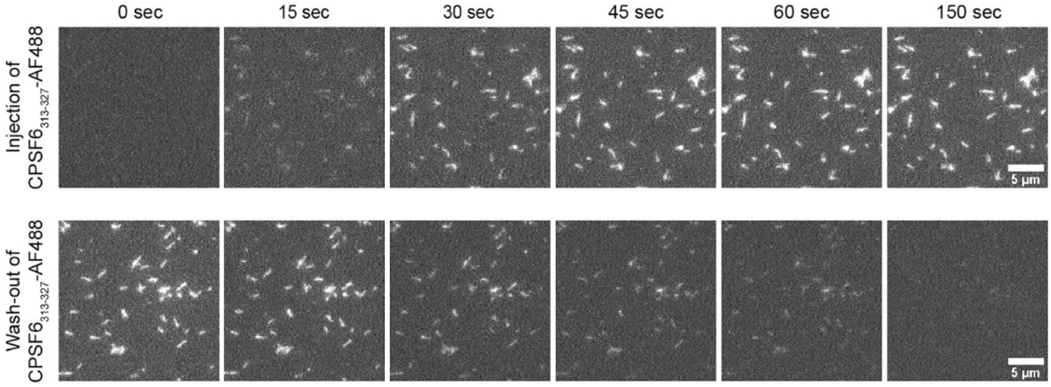


**Supplementary Figure S7**. **TIRF images of CPSF6$_{313-327}$-AF488 binding and dissociation.** Snapshots from the time series recorded in the CPSF6$_{313-327}$-AF488 channel during analyte injection and wash-out are shown at the top and the bottom, respectively.

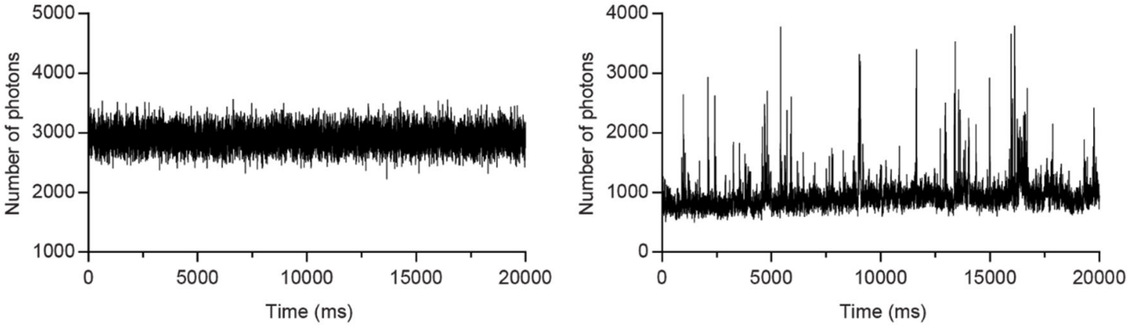


**Supplementary Figure S8. Single molecule spectroscopy of proteins produced by cell-free expression using *Leishmania* extract.** The GFP trace (left) is characteristic of the diffusion of a monomeric protein[5] while the GFP-CPSF6 trace (right) shows large photon bursts indicating the presence of higher order oligomers in solution.
